# REVERSE: A user-friendly web server for analyzing next-generation sequencing data from *in vitro* selection/evolution experiments

**DOI:** 10.1101/2022.03.21.485196

**Authors:** Zoe Weiss, Saurja DasGupta

## Abstract

Next-generation sequencing (NGS) enables the identification of functional nucleic acid sequences from *in vitro* selection/evolution experiments and illuminates the evolutionary process at single nucleotide resolution. However, analyzing the vast output from NGS can be daunting, especially with limited programming skills. No single platform exists that performs all the steps necessary to generate publishable results starting with raw sequence data. We developed REVERSE (Rapid EValuation of Experimental RNA Selection/Evolution) (https://www.reverseserver.org/), a web server that incorporates an integrated computational pipeline through a graphical user interface, which performs both pre-processing and sequence level analyses within minutes. FASTQ files from multiple rounds are quality filtered, dereplicated, and trimmed before being analyzed by two pipelines. The first pipeline counts, sorts, and tracks enrichment of unique sequences and tracks the enrichment of sequence motifs. It also identifies mutational intermediates present in the sequence data that connect two input sequences. The second pipeline sorts similar sequences into clusters and tracks enrichment of peak sequences. It also performs nucleotide conservation analysis on the cluster of choice. Both pipelines generate downloadable high-resolution figures. Collectively, REVERSE is a one stop-solution for the rapid analysis of NGS data obtained from *in vitro* selection/evolution experiments that obviates the need for computational expertise.

## INTRODUCTION

*In vitro* selection/evolution is a powerful technique to isolate nucleic acids with desired functions without a priori knowledge of their sequences by imposing selection pressure upon combinatorial libraries (1-3). Functional nucleic acids such as aptamers and (deoxy)ribozymes identified by this technique have been used in diagnostics and therapy (4-6), and *in vitro* evolution has been used to study the intricacies of molecular evolution (7-14) and explore the biochemical capabilities of nucleic acids in the context of the origin of life (15-17). With the advent of next generation sequencing (NGS), it has been possible to investigate the outcomes of combinatorial selections in unprecedented detail as NGS provides expansive sequence coverage. In addition to identifying hundreds of thousands of sequences that possess the desired phenotype, analysis of NGS data from each round may reveal the evolutionary trajectory of each sequence under changing selection pressures, thereby generating a high-resolution picture of the evolutionary process.

Sequence files obtained from NGS are usually too large to be manipulated without the use of the command line or specialized software. As a result, it is practically impossible to extract the most basic, yet essential information needed to biochemically validate selection outcomes without considerable bioinformatics expertise. NGS data must first be pre-processed to filter out ‘low quality’ sequences and remove sequencing adapters and constant regions (such as primer binding sites). Next, identical sequences must be binned (dereplication) in order to identify and count all the different sequences present in the data (unique sequences) and converted to their reverse complements, if they are from reverse reads. Usually, an arbitrary subset of the most abundant sequences is chosen for biochemical characterization. However, in order to sample the diversity of selected sequences, it is beneficial to cluster closely related sequences and characterize the most abundant sequence from each cluster (referred to as peak sequence). NGS data from all rounds of selection can be used to quantify sequence enrichment, investigate evolutionary fitness landscapes, and identify nucleotides important for function through nucleotide conservation analysis. However, these results are inaccessible to experimentalists without significant programming experience. Therefore, the field has largely depended on custom computational pipelines created by individual labs through collaborations with bioinformaticians. The unavailability of a standardized pipeline to analyze NGS data from *in vitro* selection/evolution experiments and more importantly, the absolute reliance on programming expertise that most experimentalists lack, has limited the wide implementation of NGS technology to combinatorial selections.

We developed REVERSE (Rapid EValuation of Experimental RNA Selection/Evolution) with the goal of democratizing bioinformatic analysis of NGS data from *in vitro* selection/evolution experiments by removing all computational barriers. REVERSE is an integrated computational pipeline that runs a set of custom scripts written in Python and interacts with the user through an intuitive web-based graphical interface. The entire pipeline requires a single upload step at the beginning where the user uploads a raw FASTQ file for each selection round. The pre-processing module generates spreadsheets containing quality filtered, trimmed, and dereplicated sequences with their read counts. Then the user is offered two pipelines – the first analyzes individual sequences and the second clusters similar sequences before analyzing sequence clusters. REVERSE identifies the most abundant sequences/peak sequences of each abundant clusters and quantifies their enrichment during selection. More advanced functionalities like identification of mutational intermediates between two sequences, sequence motif search, or conservation analysis are also available. In addition to spreadsheets, REVERSE outputs downloadable publication quality figures for most results. To demonstrate its utility, we applied the REVERSE workflow to a subset of NGS data (four rounds) obtained from a previous *in vitro* selection experiment designed to isolate ligase ribozymes (18). The entire REVERSE pipeline to analyze 100,000 sequences per round took about 23 minutes to complete, which includes about 4 minutes for file upload and pre-processing. In addition to analyzing outcomes from nucleic acid selection/evolution, REVERSE may be used for the computational needs of any experiment that involves combinatorial libraries such as peptide and protein selection using mRNA (19) or phage display (20) and high-throughput mutational analysis of (deoxy)ribozymes (21). Therefore, REVERSE represents the first web-based computational platform for rapid analysis of NGS data acquired from combinatorial selections that enables users to take full advantage of their experimental results without having to write a single line of code.

## MATERIALS AND METHODS

### Overview

The *Analyze* tab is the gateway to the REVERSE workflow where sequence data files are pre-processed and analyzed in two separate pipelines – the first pipeline analyzes individual sequences and the second pipeline clusters sequences before analyzing sequence clusters (Figure 1). REVERSE takes as input one zipped FASTQ file (e.g., ‘filename.fastq.zip’) per round and does not merge paired-end reads from forward and reverse runs. As most combinatorial libraries are <100 nt, single-end reads are usually sufficient to cover the library sequence. REVERSE offers two types of outputs – spreadsheets (CSV files) and downloadable visualizations. Spreadsheets contain lists of sequences with their respective read counts after pre-processing steps such as quality filtering and/or trimming. Spreadsheets allow easy manipulation of sequence data and importantly, enables instant identification of selected sequences for further biochemical validation. Visualizations include high resolution line plots, 3D bar plots, heatmaps, and selection statistics tables, that may be downloaded from the web page. All visualizations are accompanied by raw data that may be used to create customized figures.

**Figure 1.**
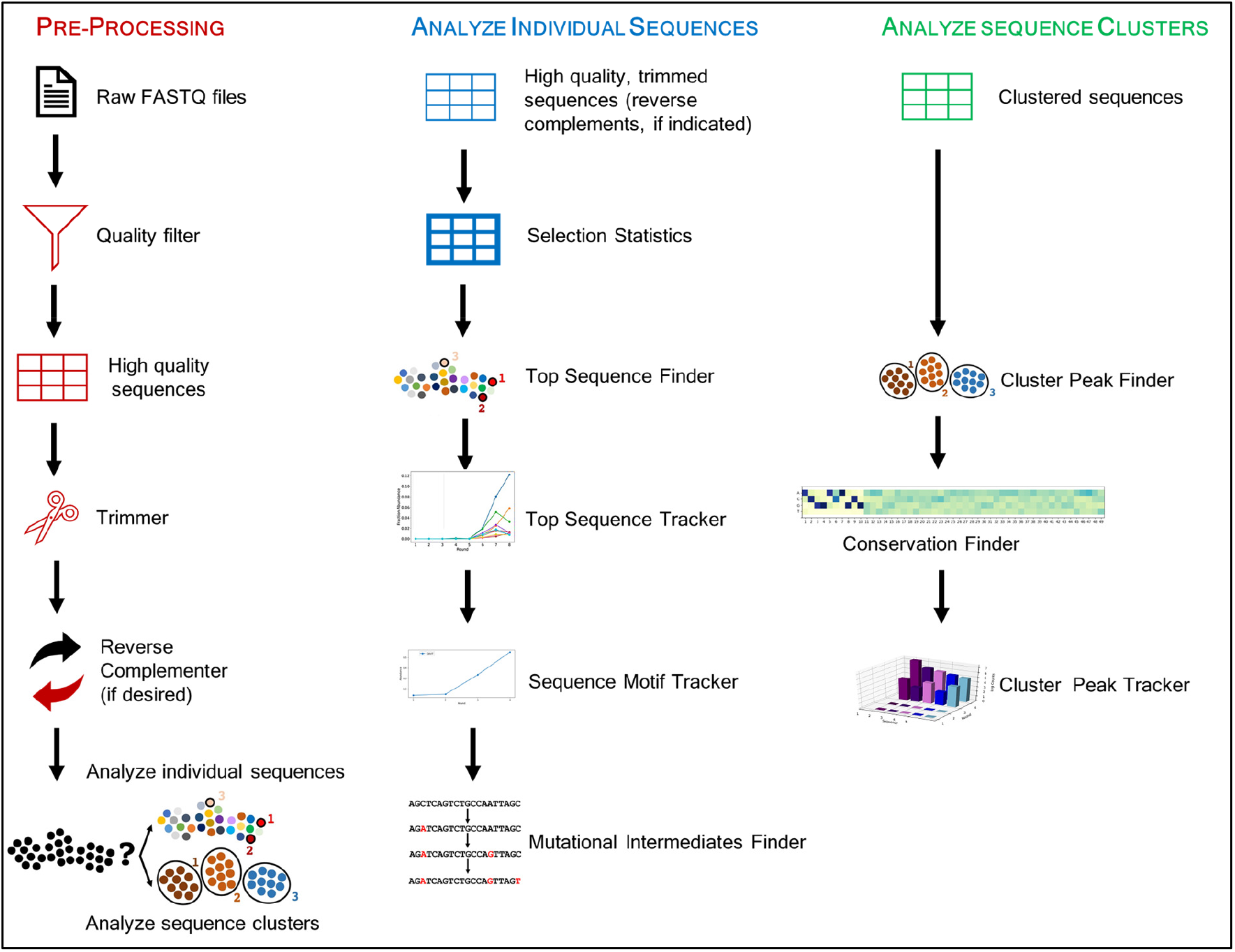
The REVERSE workflow: from raw next-generation sequencing data to data visualization with a few clicks. REVERSE pre-processes raw FASTQ files from combinatorial selections and rapidly identifies the most abundant sequences. It clusters similar sequences and identifies the most abundant clusters, tracks the enrichment of individual sequences, cluster peak sequences, and user-defined sequence motifs. REVERSE also performs covariation analysis on sequence clusters and searches sequence data for intermediates between two user-defined sequences.

### Pre-processing module: Filtering by quality, dereplication, trimming, and batch conversion to reverse complements, if required

The first page of the *Analyze* tab contains the first four steps of the pre-processing module that determine the operations to be performed on the sequence data files, once uploaded (Figure 2). *Step 1* allows the user to input the number of files they want analyzed (usually corresponds to number of rounds). REVERSE can analyze a maximum of ten files at once; however, computational time is reduced with fewer files. For experiments with more than ten rounds, users may analyze data from representative rounds that capture functional enrichment or analyze every other round, if possible. Sequencing results from replicate experiments or from parallel selections performed under different selection conditions, can also be analyzed via the multiple file upload functionality of REVERSE.

**Figure 2.**
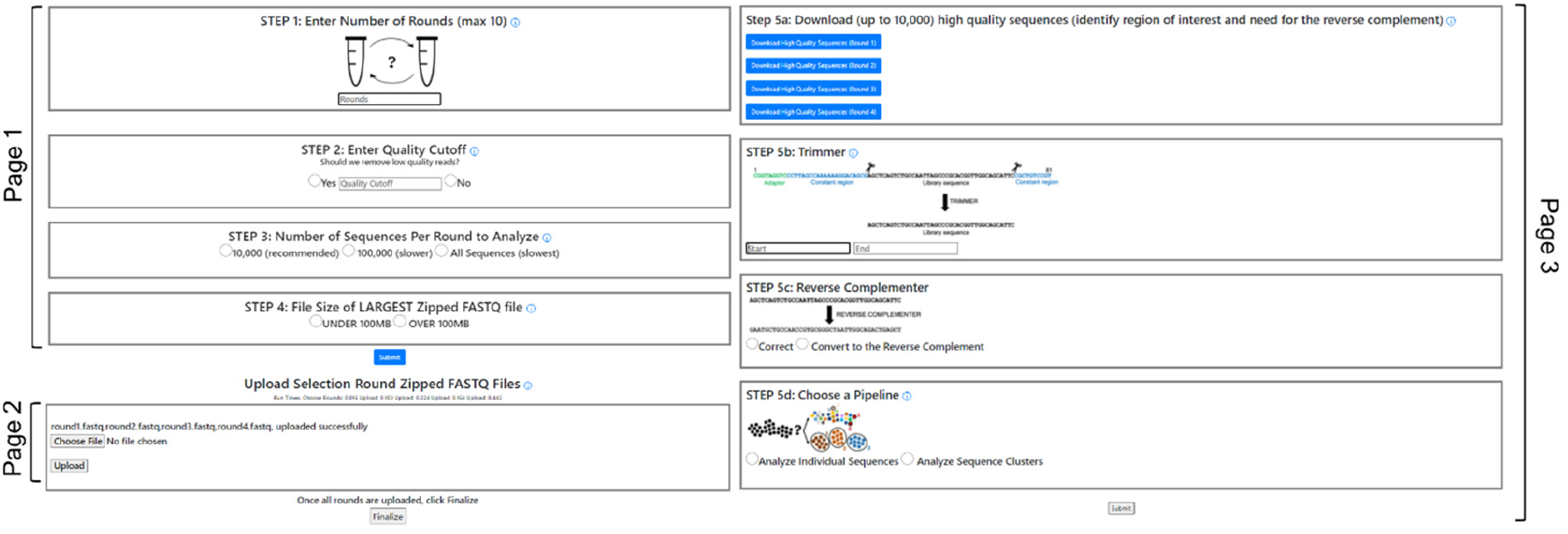
The pre-processing module of REVERSE. Page 1 sets up the parameters for subsequent analysis, page 2 provides the uploading interface for compressed FASTQ files, and page 3 displays the actual pre-processing tools. The pre-processing module bins identical sequences (dereplication) and calculates the number of reads for each dereplicated sequence. Step 5a provides a list of dereplicated sequences, the Trimmer tool in Step 5b removes undesired regions from these sequences, and the Reverse Complementer tool in Step 5c converts sequences to their reverse complements if the user desires. This tool is useful for analyzing reverse reads. Finally, Step 5d offers the user two pipelines of the analysis module of REVERSE.

*Step 2* reads the base calling accuracy at each nucleotide position (Q score) of every sequence and filters out sequences that have base calling errors larger than the user-defined cutoff. For example, a cutoff input of 98% will filter out sequences that have >2% of their nucleotides with a Q score < 20 (i.e., >1% error in base calling). *Step 3* accepts the number of sequences in each file the user wants analyzed, which could be the first 10,000, the first 100,000, or all sequences. Analyzing a subset of the sequence data lowers processing times, while preserving important information such as the identities of the most abundant sequences and their relative abundances and enrichment during selection (see Results). In *Step 4*, the user enters the size of the largest input ZIP file to determine the upload strategy. Most raw sequence files from *in vitro* selection/evolution experiments are under 100 MB when compressed; however, files larger than 100 MB can be accommodated in REVERSE with an extra processing step. The second page of the *Analyze* tab permits the user to upload multiple FASTQ files to be analyzed simultaneously. Files under 100 MB can be directly uploaded here (Figure 2). For files over 100 MB, both Mac and Windows users may use the Galaxy server (https://usegalaxy.org/) to filter out low quality sequences first and use a compressed form of this smaller output file as input for REVERSE. Alternatively, Mac users may download a simple bash script to split large files. This script splits and downloads these files to the user’s desktop. For example, this script will generate two smaller files, ‘roundXaa’ and ‘roundXab’ from ‘roundX’, a single >100 MB file. The user can then upload these smaller files in the *Upload Split Zipped FASTQ Files* page, where REVERSE will merge the two files. The commands needed for this operation are provided in this page.

Once all files are uploaded, REVERSE removes sequences with quality scores below the user-defined cutoff, bins identical sequences and counts the number of times each unique sequence was read for each file. *Step 5a* gives the user the option to view the ‘high-quality’ sequences for each uploaded sequence file in downloadable CSV files (Figure 2). This is particularly helpful in identifying the region of interest in each sequence, so that flanking constant regions such as barcode sequences and primer binding sequences can be removed. *Step 5b* allows the user to define the boundary of the region of interest by entering its first and last nucleotides in the *Trimmer* tool. In *Step 5c*, the user indicates if the uploaded files are from forward or reverse sequencing runs. The *Reverse Complementer* tool perform batch conversion of all sequences derived from reverse sequencing runs. Finally, in *Step 5d*, the user selects one of two pipelines of the analysis module of REVERSE. While the *Analyze Individual Sequences* pipeline performs a series of computations on all dereplicated sequences, the *Analyze Sequence Clusters* pipeline first clusters closely related dereplicated sequences before analyzing the most abundant sequence clusters.

### Analysis module: Analyze Individual Sequences

The *Analyze Individual Sequences* pipeline consists of the following six tools (Figure 3):

**Figure 3.**
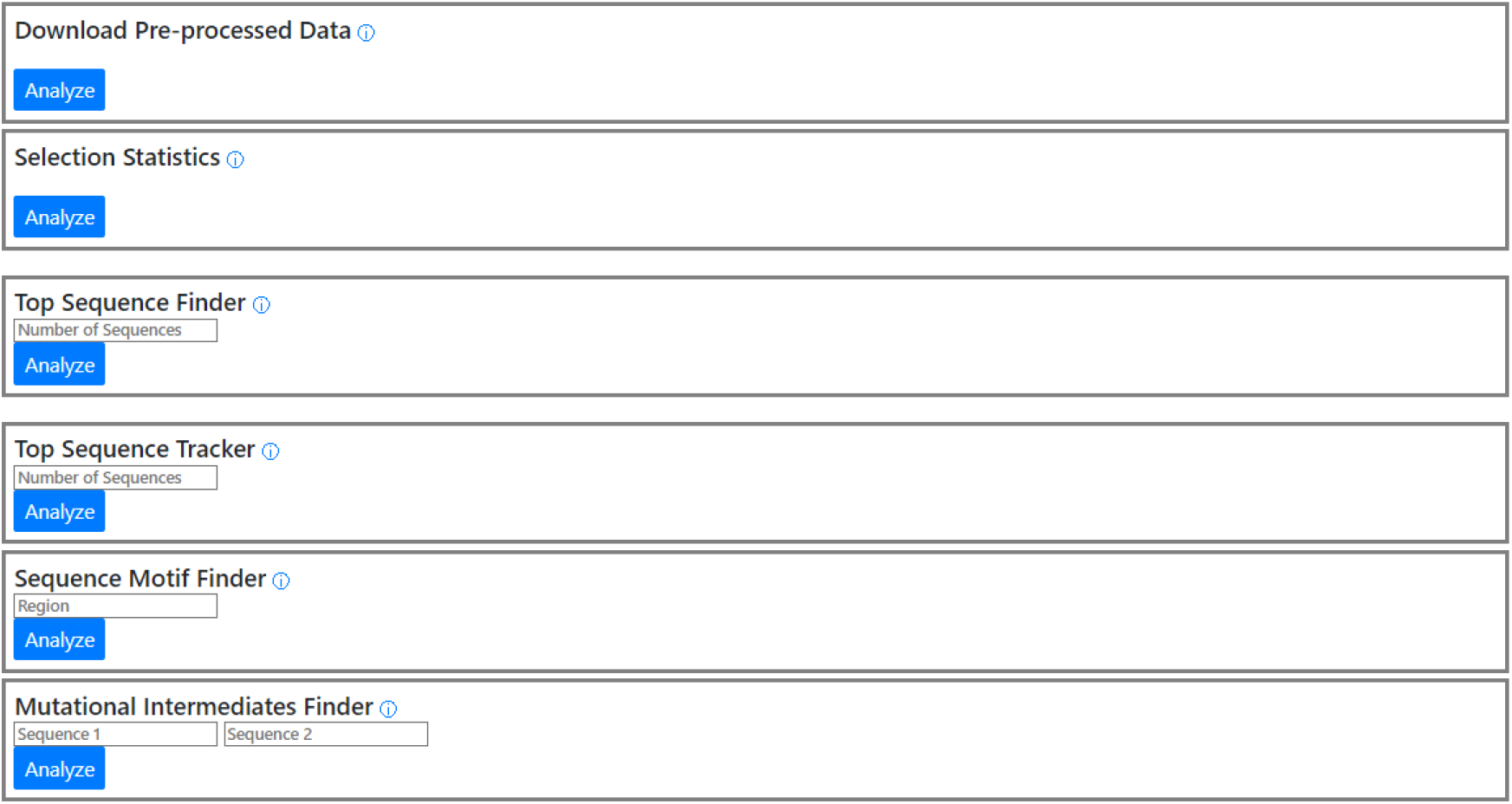
The *Analyze Individual Sequences* pipeline of the analysis module of REVERSE. This pipeline allows the user to download pre-processed data in easy-to-use spreadsheets that contains a list of unique sequences sorted according to their abundances, accompanied by their read counts. The Selection Statistics tool generates a table summarizing the selection outcome. The Top Sequence Finder tool identifies the *N* (user-defined) most abundant sequences in the last round data and the Sequence Tracker tracks the enrichment of the abundant sequences across rounds. The Sequence Motif Finder tool searches all uploaded sequence files for a user-defined nucleotide string and outputs a spreadsheet for each round that contains only sequences with the desired nucleotide string. It also generates a graphic illustrating the enrichment of such sequences across rounds. The Mutational Intermediates Finder tool searches sequence files to discover the shortest mutational path between two given sequences.

1. *Download Pre-Processed Data* outputs a CSV file, for each input FASTQ file, containing quality filtered sequences that are trimmed to the desired length, and, if desired, converted to their reverse complements. Each output file contains up to 10,000 dereplicated sequences that are sorted according to their read counts. This tool provides rapid access to all sequences isolated from selection and enables the user to inspect and manipulate sequencing data in a spreadsheet. This is one of the most sought-after results in most combinatorial selections.
2. *Selection Statistics* outputs a table containing the number of total sequences, number of high-quality sequences, number of unique sequences, and percent unique sequences for each round. This tool may be used to evaluate selection outcomes. A decrease in the fraction of unique sequences, indicates sequence enrichment, which is usually accompanied by functional enrichment and could point to a successful selection.
3. *Top Sequence Finder* outputs the *N* most abundant sequences from the final round of selection with their read counts, where *N* is user-defined. This tool allows identification of the subset of abundant sequences the user plans to characterize biochemically. These sequences are often the most active (i.e., possess the highest fitness) for the desired function.
4. *Top Sequence Tracker* plots the fractional abundance of each of the *N* most abundant sequences against selection round. In addition to visualizing sequence-level population dynamics of selection/evolution across rounds, this tool visually depicts the point in the selection where enrichment becomes evident. The tool can therefore be used to monitor selection progress even before the desired activity is detected experimentally in selection pools. Since this tool compares fractional abundances of the most abundant sequences across sequence files, it can be used to test selection reproducibility (where files represent experimental replicates instead of outputs from different rounds) or analyze the differences in sequence enrichment under diverse selection pressures (where each file represents a different selection condition).
5. *Sequence Motif Finder* outputs a spreadsheet for each round containing only sequences that possess the user-defined motif and plots the fractional abundances of these sequences across rounds. A sequence motif in this context is defined as a continuous string of nucleotides and is distinct from a structural motif such as a pseudoknot or a stem loop, which can be formed by divergent sequences. Sequence motifs must be defined using standard nomenclature – specific nucleotides: A, T, G, C; purines: R, pyrimidines: Y, and any nucleotide: N. Functional enrichment during selection is often accompanied by the enrichment of sequences that possess specific sequence motifs. These sequence motifs may constitute structural features important for activity and/or directly interact with the substrate (in case of (deoxy)ribozyme selections).
6. *Mutational Intermediates Finder* takes as input two sequences and performs an advanced search across all sequence files to identify the shortest mutational path between the two. Paths may comprise of sequences that differ from each other by one or more point mutations depending on the extent of sequence space coverage in the specific experiment. This tool can be used to discover neutral or almost-neutral mutational pathways between any two sequences. With experiments that enable complete or near-complete coverage of sequence space (11) *Mutational Intermediates Finder* can potentially identify neutral networks between distinct functional RNAs. The existence of neutral networks between functional RNAs such as ribozymes has significant implications for the evolution of RNA-based early life (22-24).

### Analysis module: Analyze Sequence Clusters

Closely related RNA sequences usually adopt similar structures and therefore can be considered to be members of the same RNA class. To identify multiple classes of functional RNAs for downstream characterization, selected sequences are often clustered through multiple sequence alignments and the most abundant sequence (peak sequence) from each of the most abundant clusters, is characterized. To address this need, we created the *Analyze Sequence Clusters* pipeline which consists of the following four tools (Figure 4):

**Figure 4.**
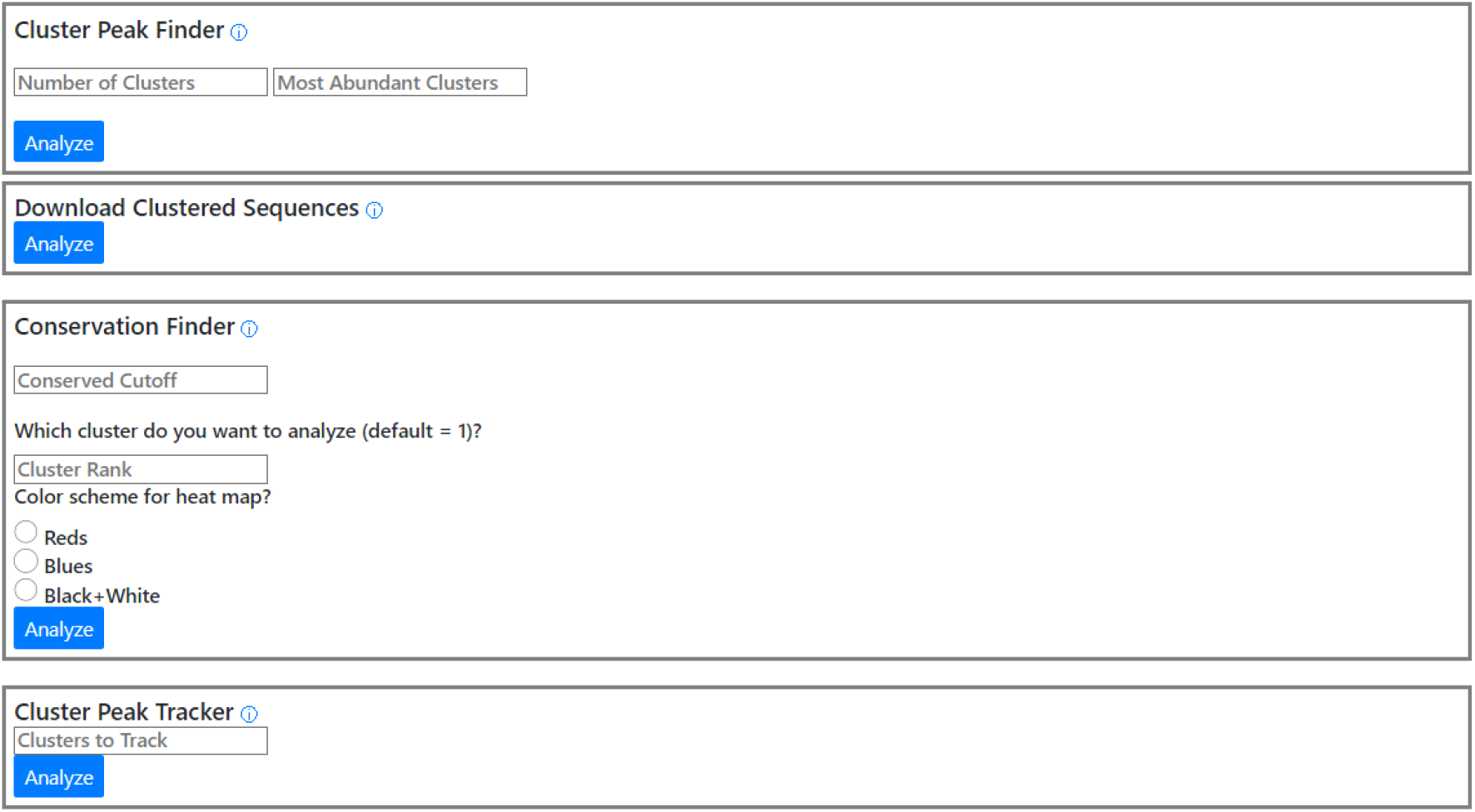
The *Analyze Sequence Cluster* pipeline of the analysis module of REVERSE. This pipeline allows the user to download a spreadsheet containing pre-processed sequence data with closely related sequences sorted into clusters. Each sequence is assigned a cluster number. Cluster Finder outputs the peak sequences (and their read counts) from the *N* (user-defined) most abundant clusters. Conservation Finder quantifies the distribution of each of the four nucleotides at each position within the cluster of choice and outputs the results in a customizable heatmap. Cluster Tracker quantified the enrichment of peak sequences of the *N* most abundant clusters.

1. *Cluster Peak Finder* outputs peak sequences (with their read counts) from *N* sequence clusters, where *N* is user-defined and cannot exceed 20. The most abundant peak sequences usually represent distinct solutions for accomplishing the selected function. This tool, therefore, provides easy access to the most abundant clusters peak sequences, which is one of most desired results from *in vitro* selection experiments.
2. *Download Clustered Sequences* allows the user to access the *N* sequence clusters from the *Cluster Peak Finder* tool. As this tool, like other clustering and multiple sequence alignment software, is time and memory intensive, currently it is unable to cluster all sequences from several large files at once. The user may perform clustering with one or two input files at once. This is not a problem for most users who would likely analyze data from only the final round of selection. The tool outputs CSV files containing up to 1000 sequences (with read counts) that are assigned to specific clusters (cluster 0 onwards). Using the Sort function in applications like Microsoft Excel, the user can organize sequences according to their cluster number and further sort sequences in a cluster according to their read counts.
3. *Conservation Finder* outputs a customizable heatmap that illustrates the distribution of nucleotides at each position within a user-specified cluster. Nucleotide variations at each position may be used to predict the importance of each nucleotide to the sequence’s fitness.
4. *Cluster Peak Tracker* outputs a plot containing the fractional abundances of the peak sequences of the *N* most abundant clusters across rounds. Like *Sequence Tracker*, this tool can evaluate selection progress before desired activity is detected in sequence pools. This tool is more useful as it reveals information about the population dynamics of distinct RNA classes.

### Implementation

The REVERSE web server is built using PythonAnywhere, an online integrated development platform running server-based Python scripts for web development. The back end of REVERSE is created in Python (https://www.python.org/downloads/release/python-3101/) with Flask (https://flask.palletsprojects.com/en/2.0.x/). The various computational tools in REVERSE are written as custom scripts in Python. Comprehensive documentation can be found in the Supplementary Data file and on the web page, which also includes a video walkthrough with demo files. The front end of REVERSE consists of a graphical user interface created with HTML and CSS scripts in PythonAnywhere. The basic layout of each page is based on designs from the Bootstrap v5.1 library (https://getbootstrap.com/). All data visualizations in REVERSE are generated using the Matplotlib and Seaborn libraries in Python. REVERSE has been tested with various combinations of Linux/ MacOS/Windows 10 and Chrome/Firefox/Edge/Safari (Supplementary Table S1).

### Comparison with similar software

REVERSE is the only software that performs both pre-processing and sequence analysis on NGS data obtained from combinatorial selections and is the only GUI-based platform to do so (Supplementary Table S2). Although software/web servers catering to RNA-seq data analysis for biological applications such as differential gene expression exist (25,26), there are next to no resources for analyzing sequence files from *in vitro* selection/evolution experiments, and therefore multiple software must be used. For example, a collection of command line software is often used to pre-process raw FASTQ files. Quality filtering may be performed using FastQC (https://www.bioinformatics.babraham.ac.uk/projects/fastqc/), constant regions may be removed using Cutadapt, cutPrimers, or Trimmomatic (27-29) and paired-end reads may be merged using PANDAseq (30) or PEAR (31). The Galaxy server provides a web-based GUI to implement some of these tools; however, it does not perform bioinformatic analysis such as dereplicating, counting, and clustering that are essential for analyzing results from *in vitro* selection experiments. EasyDIVER (32), a command line software, can perform paired-end joining, dereplication, and trimming, but performs no further bioinformatic analysis. FASTAptamer (33) is currently the only software developed to specifically analyze NGS data from combinatorial selections. FASTAptamer counts dereplicated sequences, compares relative abundances in sequences present in two files, searches for degenerate sequence motifs and performs sequence clustering. However, FASTAptamer does not contain any pre-processing functionalities and its command line interface limits wide adoption by experimentalists not familiar with the command line. Further, FASTAptamer primarily outputs results within the command line and does not generate easily usable sequence lists or visualizations.

The REVERSE server is a significant advance in the field of combinatorial selection, because not only is it a one-stop solution to its most common bioinformatic needs, but also the only resource that interacts with users using a GUI and requires no coding expertise or any familiarity with the command line. The computational pipelines in the Galaxy server or the FASTAptamer toolkit are hierarchical where output files from previous steps are inputs for the next steps. This requires multiple input files to be properly processed before moving forward in the pipeline. REVERSE overcomes this limitation by implementing custom scripts that run in the background, which allows users to upload multiple files in single step at the beginning and freely use the tool of their choice without being constrained by prior steps. In addition, REVERSE is the only software to output downloadable publication quality visualizations by analyzing NGS data from combinatorial selections. Therefore, by simply uploading the raw FASTQ files obtained from sequencing and entering the required parameters in each step, users will be able to access the most important results from selection experiments, which will accelerate further experimental characterization and the provide publishable figures to accompany these results.

## RESULTS

### Case Study: The REVERSE Workflow

To illustrate the various functionalities of REVERSE, we analyzed four rounds (two before functional enrichment was visible and two after) of raw NGS data from a previous *in vitro* selection experiment designed to isolate ribozymes that catalyze templated ligation of RNA oligomers activated with 2-aminoimidazole (18). Users may try out all the tools REVERSE has to offer in the *Tutorial* tab with truncated versions of these files (demo files). After entering pre-processing parameters in *Steps 1-4* like number of rounds (4), quality cutoff, (98) number of sequences to be analyzed for each file (10,000) and file size (under 100 MB), four FASTQ files corresponding to reverse reads from each of the four rounds were uploaded to the pre-processing module. After inspecting high-quality sequences in downloaded spreadsheets (one for each round) in *Step 5a*, the region of interest (representing the 40 nt randomized library) was identified between nucleotides 32 and 71. This information was used to remove extraneous sequences in *Step 5b*. Since the uploaded sequence data was obtained from reverse reads, all sequences were converted to their reverse complements in *Step 5c*. In *Step 5d*, the *Analyze Individual Sequences* pipeline was selected first.

We downloaded pre-processed data for each round as separate spreadsheets, which allowed us to readily identify the most abundant sequences from each round. Selection outcomes from the *Selection Statistics* tool revealed a marked decrease in the fraction of unique sequences (from 0.997 in round1 to 0.246 in round 4) indicating enrichment (Figure 5A). The *Top Sequence Finder* and *Top Sequence Tracker* tools were used to output the five most abundant sequences in the round 4 sequence data and track its fractional abundance through rounds 1-4 (Figure 5B, C). The visualization shows that significant enrichment of abundant sequences occurred in round 3. Next, we searched for the ‘TGCGG’ motif in the sequence data. Prior biochemical results suggested that this motif likely helps these ligase ribozymes bind their RNA substrate via complementary base-pairing (18). The *Sequence motif tracker* tool generated spreadsheets containing sequences that possess this motif and a plot illustrating how these sequences were enriched across selection. Marked enrichment was observed in round 3, which coincides with the enrichment of the most abundant sequences (Figure 5C, D), underscoring the importance of this motif in ribozyme function. The test dataset was not suited for the *Mutational Intermediates Finder* tool and therefore this tool was not tested here.

**Figure 5.**
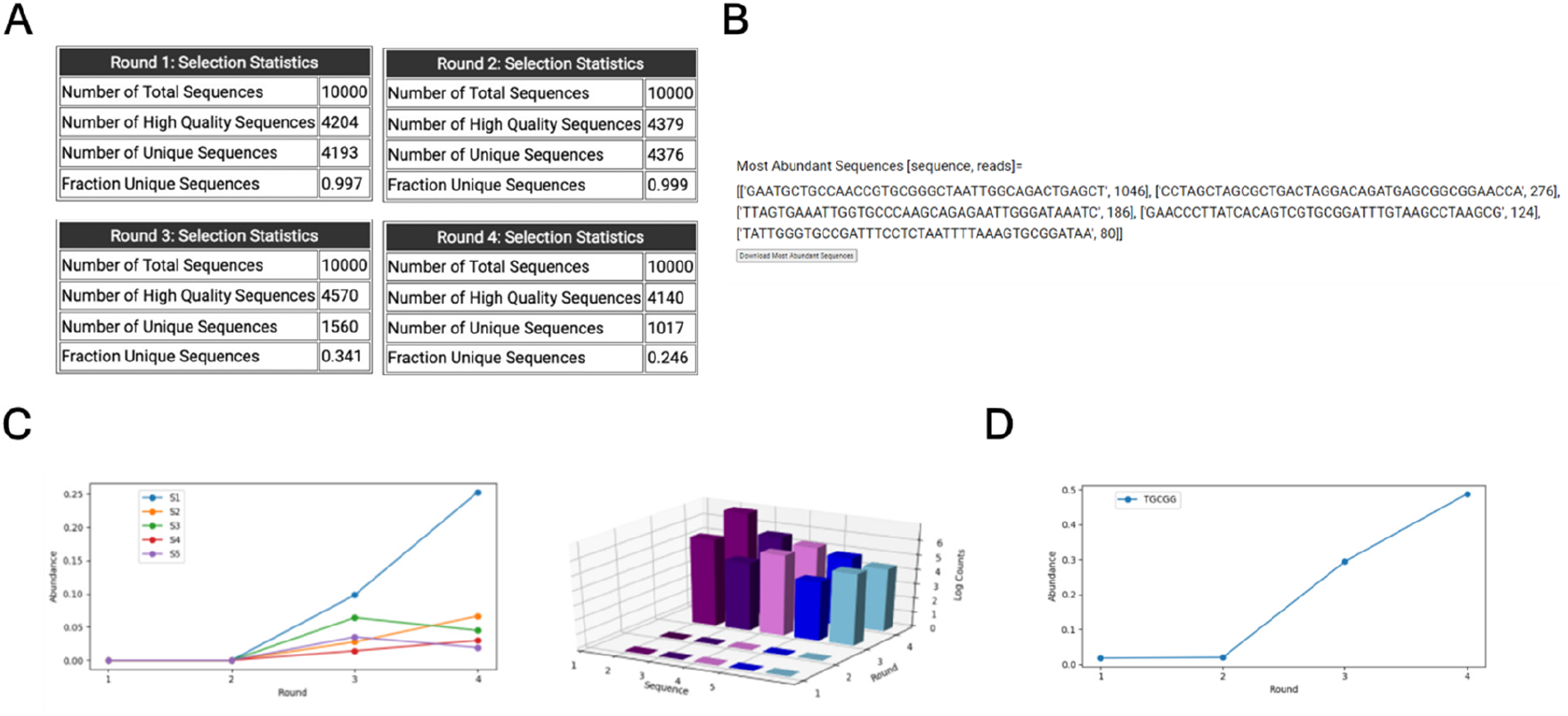
The *Analyze Individual Sequences* pipeline: **A**. summarizes selection outcomes; **B**. identifies the most abundant sequences; and **C**. plots their relative abundances across selection. It also **D**. lists all sequences containing a user-defined sequence motif and graphically depicts their enrichment.

After completing the first pipeline, we returned to the last page of the pre-processing module (*Step 5*), and selected the *Analyze Sequence Clusters* pipeline in *Step 5d* after re-entering input parameters in *Steps 5b and c*. This pipeline allows immediate access to spreadsheets with sequence clusters from each round and the identities of their peak sequences (Figure 6A, B) after taking as input, the maximum number clusters. To test the *Conservation Finder* tool, we analyzed the most abundant cluster with a conservation cutoff of 98%. Certain positions like 6, 10, 20, 25, and 37 were found to be highly conserved, while other positions like 4, 23, 38 were less conserved (Figure 6C). Finally, the *Top Cluster Tracker* tool was used to quantify the relative abundances of the five most abundant peak sequences along the selection trajectory (Figure 6D).

**Figure 6.**
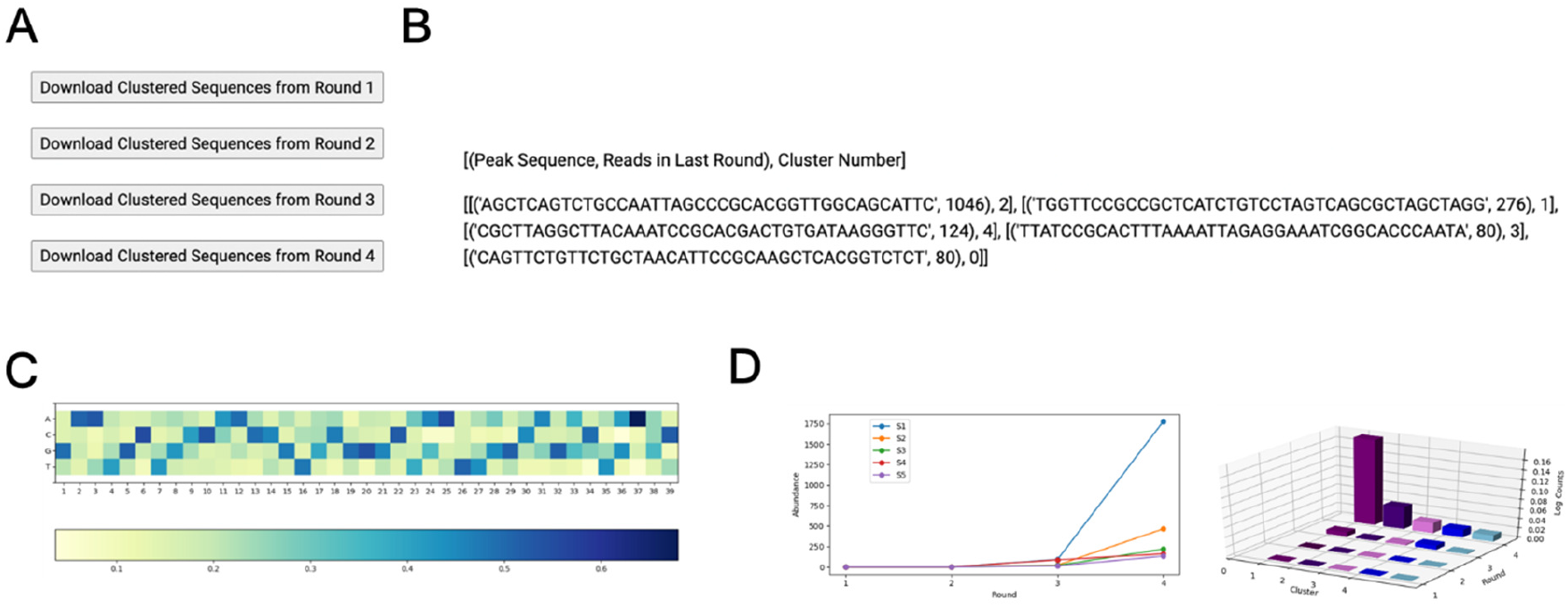
The *Analyze Sequence Clusters* pipeline: **A**. outputs clustered sequences from each round; *B*. identifies the peak sequences in each of the most abundant clusters; **C**. quantifies nucleotide conservation at each position and depicts the results in a customizable heatmap and **D**. plots the enrichment of peak sequences across selection.

### Case Study: Computation speed evaluation

We measured the computation time for each tool with either 10,000 or 100,000 sequences per sequence file (on Chrome browser). For 10,000 sequences, file upload and pre-processing steps were completed in 2 minutes, whereas it took 4 minutes for 100,000 sequences. The *Analyze Individual Sequences* and *Analyze Sequence Clusters* pipelines took 40 and 170 seconds for 10,000 sequences, respectively. For 100,000 sequences, these pipelines took 6.5 and 12 minutes, respectively. Additionally, we timed each step when REVERSE was used to analyze all sequences from a single file that had been quality filtered by the Galaxy server. The pre-processing module (without quality filter) took 1.2 minutes, and the *Analyze Individual Sequences* and *Analyze Sequence Clusters* pipelines took 1.5 and 7.5 minutes, respectively. A detailed breakdown for each tool in each instance is provided in Supplementary Table S3.

### Case Study: Applicability of analyzing a subset of sequence data

As analyzing a subset of the total number of sequences significantly reduced computation times, we wondered if results obtained from the analysis of 10,000 sequences could be extrapolated to the analysis of more sequences. We compared results in each case using the sequence file from the final round of selection (Supplementary Table S4). The fraction of unique sequences, which indicates enrichment, were somewhat comparable between 10,000 (0.246) and 100,000 (0.148) sequences. The most abundant sequence was identical in all three cases and their relative abundances were nearly identical. These results suggest that analyzing a subset of the sequence data may provide similar outcomes while significantly accelerating the process.

## DISCUSSION

We developed the REVERSE web server as a one-stop solution for the bioinformatic needs of analyzing next-generation sequencing data obtained from combinatorial selections. REVERSE is completely GUI-based, which allows experimentalists without programming skills or familiarity with the command line to integrate NGS with their combinatorial selection experiments. To our knowledge REVERSE is the only computational platform for NGS data analysis of *in vitro* selection experiments that contains an integrated pre-processing module. The REVERSE workflow is intuitive and streamlined, where the user uploads multiple raw FASTQ files once without the need to use processed files for subsequent steps. REVERSE identifies the most abundant sequences/sequence clusters, graphically tracks the relative abundances of selected sequences across rounds of selection and quantifies sequence enrichment in minutes. By removing the need for writing custom codes and/or having to rely on bioinformatic collaborations, we think REVERSE will drastically reduce the time lag between NGS data acquisition and biochemical characterization of isolated sequences. In addition to generating the results common to most combinatorial selections, REVERSE performs more advanced computations such as conservation analysis and outlining mutational paths between source and target sequences. Although REVERSE was developed for analyzing nucleic acid selection/evolution experiments, it can be used to analyze data generated by other experiments that use combinatorial libraries.

Web-based bioinformatic platforms for analyzing large NGS data files require non-trivial computational resources and are time-intensive. By allowing users to analyze a representative subset of their data, REVERSE can significantly reduce analysis times. For example, the pre-processing module is two-fold faster for the same dataset if 10,000 sequences is analyzed instead of 100,000 (Supplementary Table S3). Although a subset of the data is likely sufficient for most combinatorial selections needs, certain tasks require greater coverage of sequence data. These include detection of enrichment in earlier rounds, conservation analysis of specific sequence clusters, discovering neutral networks between two distinct RNAs, and broader investigations into the nature of evolutionary fitness landscapes. The current memory constraints stem from the ‘3 GB per process’ RAM limit imposed by PythonAnywhere, therefore, REVERSE cannot analyze all sequences from multiple sequence files simultaneously. With increasing traffic, we plan to host REVERSE in a private server within the PythonAnywhere cluster which will remove the current memory limits. Presently, if users wish to analyze all sequences (and not a subset), they may upload pre-processed files from the Galaxy server to reduce the number of sequences in each file and consequently reduce computational demands. Alternatively, users may analyze sequence data from the final round if their primary goal is to identify the most abundant sequences.

We are also working on implementing computational strategies to fragment sequence files into readily analyzable blocks (∼100,000 sequences), which will undergo parallel pre-processing before being concatenated for the analysis module (34). This strategy will also be utilized for tools in the analysis module that benefit from greater sequence coverage. Once REVERSE is able to handle multiple full-length files, we will incorporate a paired-end read merger tool, which will reduce sequencing errors and thus increase high-quality sequence reads. Following this, the Trimmer tool will be updated to allow the user to isolate the region of interest by defining the identities of its flanking sequences, instead of defining the ‘start’ and ‘end’ nucleotides. This update will accommodate sequences containing indels generated due to mutations during amplification. In future iterations, we also plan to integrate the Infernal (35) secondary structure motif search to complement REVERSE’s existing sequence motif search tool and incorporate R-scape for covariation analysis (36). Since NGS can identify multiple variants of peak sequences within a cluster, multiple sequence alignments can be used to search for recurring structural motifs using Infernal and generate secondary structure models by covariation analysis using R-scape. We hope that REVERSE will allow nonbioinformatician-experimentalists to benefit from NGS technologies in their *in vitro* selection/evolution experiments.

## AVAILABILITY

REVERSE is freely available at https://www.reverseserver.org/ and does not have a login requirement. The code for REVERSE is open source and available in the GitHub repository (https://github.com/szostaklab/REVERSE).

## SUPPLEMENTARY DATA

Supplementary Data are available at NAR online.

## ACKNOWLEDGEMENT

We thank Dr. Daniel Duzdevich for helpful comments on this project and thank Dr. Jack Szostak for valuable suggestions throughout this project, critical reading of the manuscript, and procuring funding.

## FUNDING

Harvard Origins of Life Initiative Summer Fellowship, Harvard PRISE Fellowship, SETI Forward [Z.W]. Simons Foundation [290363FY18] and MGH internal funding [J.W.S.] J.W.S. is an investigator of the Howard Hughes Medical Institute. Funding for open access charge: MGH internal funding.

## CONFLICT OF INTEREST

None declared

## Notes

https://github.com/szostaklab/REVERSE

### Competing Interest Statement

The authors have declared no competing interest.

