## Supplementary data for "REVERSE: A user-friendly web server for analyzing next-generation sequencing data from *in vitro* selection/evolution experiments"

**Supplementary Material**

Zoe Weiss^1^ and Saurja DasGupta^2,3,4,*^

^1^ Department of Chemistry and Chemical Biology, Harvard University, Cambridge, MA 02138, USA.

^2^ Department of Molecular Biology, Center for Computational and Integrative Biology, Massachusetts General Hospital, Boston, MA 02114, USA.

^3^ Howard Hughes Medical Institute, Massachusetts General Hospital, Boston, MA 02114, USA.

^4^ Department of Genetics, Harvard Medical School, Boston, MA 02115, USA.

**Detailed description of each computational tool in REVERSE**

**The Pre-processing module**

***Page 1: Enter Data Parameters***

*Step 1: Enter Number of Rounds (max 10)*

A short answer text form prompts the user to enter the number of rounds (the number of sequence files they want to analyze). This answer is stored in a global variable and is referenced on Page 2 during file upload. If the user leaves the field blank, enters a non-numeric value, or a value greater than 10, the server will output an error message once the form is submitted.

*Step 2: Enter Quality Cutoff*

This form asks the user if REVERSE should remove low-quality sequences from their dataset. If they select ‘Yes’, they must enter their quality cutoff. If neither option is selected or ‘Yes’ is selected but no cutoff is entered the server will output an error message upon submission. The response to Step 2 is stored in a global variable and used on Page 3 when the algorithm pre-processes the uploaded data.

*Step 3: Number of Sequences Per Round to Analyze*

The user is asked how many sequences in the sequence data they would like REVERSE to analyze. They are provided with three options: ‘10,000 (recommended)’, ‘100,000 (slower)’, and ‘All Sequences (slowest).’ The user’s choice is stored in a global variable and is referenced on Page 2 when the algorithm uploads the selected number of sequences.

*Step 4: File Size of LARGEST Zipped FASTQ file.*

The user is prompted to select whether the size of their largest zipped FASTQ file is above or below 100 MB. Python Anywhere limits the maximum file upload size, but most data files from *in vitro* selection/evolution experiments exceed this limit. Therefore, we developed alternatives to first decrease file size through compression or to upload a compressed file in parts if they exceed 100 MB while compressed. The file size selection is stored in a global variable and dictates the instructions shown on the subsequent page.

***Page 2: Upload Data Files***

The user is prompted to upload their sequence file(s) one at a time. For a single file upload, the script unzips the file using a built-in command and deletes the original zipped file. For multiple file uploads, the script first uploads each file and then sequentially unzips them. Next, the script opens each file and reads the encoded information in list form. The script reads only the number of lines (sequences) specified in Step 3. Uploaded data are compiled into a master list to be pre-processed using inputs on Page 3.

***Page 3: Preprocess Data***

*Step 5a: Download High quality sequences.* For each uploaded file, a ‘Download High Quality Sequences’ button is displayed, prompting the user with the option to download lists of quality-filtered sequences. A click triggers a script that iterates through all sequences, removing those in which the percentage of positions with greater than one percent error exceeds the user-defined cutoff. The script then creates a CSV file where each line contains a high-quality sequence and its read count and downloads it to the user’s computer.

*STEP 5b: Trimmer*

The user is prompted to enter the start and end positions of the region of interest in each sequence. These positions are saved in global variables, and upon submission of the Page 3 form, each sequence string is truncated to include only the region bounded by the input positions. If the user fails to enter the start or end positions, enters non-numeric values, or if the start position exceeds the end position, an error message is displayed.

*STEP 5c: Reverse Complementer*

The user is given an option to choose batch conversion of all sequences to their reverse complements if their sequence data is derived from reverse reads. Their selection is saved in a global variable. This tool employs the reverse complement function from the Biopython library. If no option is selected, an error message is shown.

*STEP 5d: Choose a Pipeline*

Finally, the user must select either of two pipelines to analyze their data. Their selection is saved in a global variable. If the user does not select an option, an error message is shown.

**The Analysis module**

***Analyze Individual Sequences***

*Download Pre-processed Data*

This tool does not need any direct input but instead uses the previously pre-processed sequences from Page 3 as an input to generate a CSV file with sequences and their read counts. Sequences are sorted from the most abundant to the least abundant. These spreadsheets can be downloaded by clicking the ‘Analyze’ button followed by the appropriate ‘Download Most Abundant Sequences’ buttons for each round.

*Selection Statistics*

This tool iterates through the uploaded data files from each round, calculating the number of raw sequences, then parsing high-quality, pre-processed sequences, and finally calculating the number of high-quality sequences for each round. Next, it counts the number of dereplicated sequences (unique) for each round. The number of unique sequences is divided by the number of high-quality sequences to calculate the fraction of unique sequences in each round. Results are displayed in a table.

*Top Sequence Finder*

The user must enter the number (*N*) of most abundant sequences they want to output. This tool

sorts pre-processed sequences by abundance and identifies the *N* most abundant sequences

from the final selection round.

*Top Sequence Tracker*

This function performs the same steps as Top Sequence Finder but additionally iterates through other sequence files and counts the number of reads of each of the top sequences identified from the final round. After repeating this process for each of the *N* most abundant sequences, it determines the fraction abundance of each sequence in each round by dividing the read count of each sequence by the number of high-quality sequences in that round. Finally, the function creates a line plot using the matplotlib library to visualize the relative abundance of each sequence across rounds.

*Sequence Motif Finder*

The user inputs a sequence string containing only 'A', 'C', 'G', 'T', 'R', 'Y', or 'N'. The use of any other letter elicits an error message. This tool creates combinations of all possible motifs from the input. For example, an input of a string with a single ‘R’ generates two strings – one with ‘G’ and one with ‘A’ in place of ‘R’. Similarly, a string with ‘Y’ is converted to two strings, each with either a ‘C’ or a ‘T’ in place of ‘Y.’ A string with ‘N’ is converted to four strings, each with one of the four nucleotides. Following this, the tool searches high-quality sequences from each round for any of the substrings. Each hit increases a counter by one. Finally, the function plots the number of substring hits in each selection round using the matplotlib library.

*Mutational Intermediates Finder*

This tool requires two input sequences, which must be contained in the CSV file output from *Download Pre-processed Data,* or an error message will be displayed. The tool first determines the positions in both sequences that contain identical nucleotides by indexing through each string. Next, it iterates through all high-quality sequences and adds to a list sequences with the same nucleotide at all the positions where the input sequences overlap. This list containing all possible intermediates is the output.

***Analyze Sequence Clusters***

*Cluster Peak Finder*

The user enters the number of clusters, *K*, that they desire to group sequences into. The script uses the built-in ‘*k*-means’ function from the scikit-learn library to cluster sequences into *K* clusters. This tool generates label assignments (starting with 0) for each unique sequence denoting its assigned cluster. The tool requires another user-defined parameter, *M*, that specifies the number of peak sequences to be analyzed. The script iterates through the list of sequences sorted by read counts until it determines the peak sequences from the *M* most abundant clusters.

*Download Clustered Sequences*

This tool lists clustered sequences from *Cluster Peak Finder* in a CSV file which is downloaded to the user’s computer.

*Conservation Finder*

This tool requires three parameters: the cutoff above which a position is considered conserved, the rank of the cluster of interest (*R*), and the color scheme for the heat map. An error message is displayed if any of these parameters are not entered or entered in an incorrect format. The script selects the sequences from the final round of selection, and by reusing *Download Clustered Sequences* and *Cluster Peak Finder* functions, the script isolates the sequences from the *R* ranked cluster. It then iterates through each position for the length of the trimmed sequences and determines the fraction distribution of each nucleotide at each position using the Counter function from the collections library. This data is plotted on a downloadable heatmap using the seaborn library with four rows and the same number of columns as the length of the sequences.

*Top Cluster Tracker*

Similar to *Top Sequence Tracker*, this tool first performs the same steps as *Cluster Peak Finder*, iterates through each previous selection round, and counts the number of reads of each of the cluster peak sequences identified from the final round. After repeating this process for each of the desired peak sequences, it determines the fraction abundance of each peak sequence in each round. This is calculated by dividing the number of counts of each peak sequence by the number of high-quality sequences in the corresponding round. Finally, the function creates a line plot using the matplotlib library to visualize the enrichment of each cluster’s peak sequence.

**Supplementary Tables**

| **OS** | **Version** | **Chrome** | **Firefox** | **Microsoft Edge** | **Safari** |
| --- | --- | --- | --- | --- | --- |
| Linux | CentOS 7 | not tested | 61.0 | n/a | n/a |
| MacOS | HighSierra | 98.0.4758.109 | 61.0 | n/a | 12.0 |
| Windows | 10 | 99.0.4844.51 | 61.0 | 42.17134.1.0 | n/a |

**Supplementary Table S1.** Compatibility of REVERSE with various operating systems and browsers.

| **Features** | **REVERSE** | **Galaxy** | **EasyDIVER** | **FASTAptamer** |
| --- | --- | --- | --- | --- |
| **General features** | | | | |
| User interface | Web-based GUI | Web-based GUI | Command line | Command line |
| Input steps | Single | Multiple | Single | Multiple |
| Input type | Single-end | Single or  Paired-end | Paired-end | Single-end |
| Output:  Sequence data | Easy to use spreadsheets | FASTQ/FASTA | Text | FASTA/Text |
| Output:  Data visualization | Downloadable high-resolution figures |  |  |  |
| **Pre-processing** | | | | |
| Quality filter | 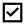 | 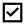 |  |  |
| Trimming | 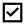 | 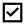 | 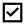 |  |
| Join paired-end reads |  | 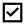 | 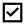 |  |
| Dereplicate and count unique sequences | 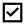 |  | 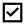 |  |
| Download sequences | 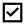 |  |  |  |
| Reverse complement | 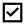 | 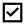 |  |  |
| **Sequence Analysis** | | | | |
| Selection statistics | 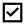  (Multiple rounds) |  |  | 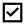  (Single round) |
| Outputs the most abundant sequences  in a spreadsheet | 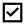 |  |  |  |
| Tracks abundant sequences during selection | 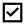 |  |  |  |
| Sequence motif search | 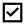  (Tracks abundance across selection) |  |  | 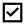  (Does not track abundance) |
| Sequence Clustering | 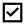  (Python algorithm) | 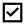 |  | 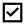  (Levenshtein edit distance) |
| Outputs the most abundant sequences  in a spreadsheet | 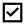 |  |  |  |
| Conservation analysis for each cluster | 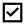 |  |  |  |
| Tracks cluster peak sequences clusters during selection | 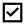 |  |  |  |

**Supplementary Table S2.** Comparison of REVERSE with related software. REVERSE is the only software that performs pre-processing and bioinformatic analysis on NGS data from *in vitro* selection/evolution experiments. The graphical interface of REVERSE allows users easy access to advanced computational pipelines that were traditionally available only to practitioners with considerable coding experience.

| **Steps** | **Four files,**  **10,000 sequences** | **Four files,**  **100,000**  **sequences** | **One file, all sequences**  **(quality filtered by Galaxy)** |
| --- | --- | --- | --- |
| **Pre-processing** |  |  |  |
| Page 1: Enter Data Parameters | 0.129 | 0.484 | 0.027 |
| Page 2: Upload Data Files | 85.303 | 79.835 | 48.91 |
| Page 3: Preprocess Data | 24.078 | 136.763 | 23.88 |
| **Analyze Individual Sequences** |  |  |  |
| Download Pre-Processed Data  (average per round) | 7 | 56 | 37.50 |
| Selection Statistics | 0.322 | 6.481 | 16.16 |
| Top Sequence Finder | 5.235 | 80.732 | 23.18 |
| Top Sequence Tracker | 6.188 | 73.95 | n/a |
| Sequence Motif Finder | 0.176 | 3.715 | 4.45 |
| Mutational Intermediates Finder | 0.062 | 0.346 | 5.58 |
| **Analyze Sequence Clusters** |  |  |  |
| Download Clustered Sequences  (average per round) | 20.584 | 72.607 | 42.56 |
| Cluster Peak Finder | 19.908 | 59.99 | 81.12 |
| Conservation Finder | 32.64 | 212.8 | 326.32 |
| Cluster Peak Tracker | 34.05 | 128.2 | n/a |

**Supplementary Table S3.** Evaluation of computation speeds (in seconds) for analyzing 10,000 or 100,000 sequences for each of the four files and all sequences for a single quality filtered file uploaded to REVERSE. Testing performs with Chrome browser.

**Supplementary Table S4.** Applicability of analyzing a subset of sequence data. The analysis module of REVERSE, when applied to a subset of the sequence data, returns comparable results.

| **Features** | **10,000 sequences** | **100,000 sequences** | **All sequences, quality filtered by Galaxy** |
| --- | --- | --- | --- |
| **Fraction of unique sequences in round 4** | 0.246 | 0.148 | 0.063 |
| **Most abundant sequence**  **(Fraction abundance)** | AGCTCAGTCTGCCAATTAGCCCGCACGGTTGGCAGCATTC  (0.23) | AGCTCAGTCTGCCAATTAGCCCGCACGGTTGGCAGCATTC  (0.23) | AGCTCAGTCTGCCAATTAGCCCGCACGGTTGGCAGCATTC  (0.21) |
